# Discovering Key Transcriptomic Regulators in Pancreatic Ductal Adenocarcinoma using Dirichlet Process Gaussian Mixture Model

**DOI:** 10.1101/2020.10.01.322768

**Authors:** Sk Md Mosaddek Hossain, Aanzil Akram Halsana, Lutfunnesa Khatun, Sumanta Ray, Anirban Mukhopadhyay

## Abstract

Pancreatic Ductal Adenocarcinoma (PDAC) is the most lethal type of pancreatic cancer (PC), late detection of which leads to its therapeutic failure. This study aims to find out key regulatory genes and their impact on the progression of the disease helping the etiology of the disease which is still largely unknown. We leverage the landmark advantages of time-series gene expression data of this disease, and thereby the identified key regulators capture the characteristics of gene activity patterns in the progression of the cancer. We have identified the key modules and predicted gene functions of top genes from the compiled gene association network (GAN). Here, we have used the natural cubic spline regression model (splineTimeR) to identify differentially expressed genes (DEG) from the PDAC microarray time-series data downloaded from gene expression omnibus (GEO). First, we have identified key transcriptomic regulators (TR) and DNA binding transcription factors (DbTF). Subsequently, the Dirichlet process and Gaussian process (DPGP) mixture model is utilized to identify the key gene modules. A variation of the partial correlation method is utilized to analyze GAN, which is followed by a process of gene function prediction from the network. Finally, a panel of key genes related to PDAC is highlighted from each of the analyses performed.

Please note: Abbreviations should be introduced at the first mention in the main text – no abbreviations lists. Suggested structure of main text (not enforced) is provided below.

## Introduction

In genetics, gene expression is one of the elementary constitutional blocks which gives rise to a phenotype from a genotype, i.e., a trait which is observable in all living cells including prokaryotes and eukaryotes. Multiple techniques are available to quantify gene expression and its regulation like DNA microarray, RNA seq, etc. The area of gene expression analysis has undergone several major advancements in biomedical research. With increased efficiency and quality, these measurements have led to improvements in disease sub-classification, gene identification problems, and studying progression characteristics of diseases ^1–7^. Biological mechanisms are dynamic in nature, therefore its activities must be supervised at multiple time points. Time-series gene expression experiments are used widely used to monitor biological processes in a time-series paradigm ^8^. Analyzing these time-series gene expression data helps in identifying transient transcriptional changes, temporal patterns, and causal effects of genes. Time-series gene expression studies can be utilized to predict phenotypic outcomes over a period of time ^9^.

DNA microarrays and RNA-Seq data have been accepted as gold standards for analyzing and measuring gene expressions across different biological circumstances ^3,5,10^. A gene is considered differentially expressed (DE) if a statistically significant difference in gene expression levels is observed between two experimental conditions. Various statistical distribution models like the Poisson and the Negative Binomial (NB) distribution are used to estimate the patterns of differential gene expression. Gene selection refers to the detection of the most significant differentially expressed genes under various conditions ^11^. Selection is done based on a combination of score cutoff and expression change threshold, which are commonly generated by the statistical design itself ^12^. Popular time course DE analysis tools include edgeR/DESeq2, TimeSeq, and Next maSigPro which are based on NB distribution model. Some DE tools like ImpulseDE2 and splineTimeR, which are based on impulse model and spline regression model between two groups respectively, are used on short time-series data ^13^.

Gene expression is a strongly regulated spatio-temporal process. Genes having identical expression patterns are associated with the same biological function. Clustering genes with similar expression pattern reduces the transcriptional response complexities by grouping genes responsible for a distinct cellular process ^14,15^. Several statistical clustering techniques have been widely used like k-means, hierarchial clustering^16,17^ and self-organizing maps^18^ to produce time-series modules. Gradually, various techniques have been developed especially for clustering time-series data. In^19^, Short Time-series Expression Miner (STEM) has been used as a clustering technique that maps genes to their representative expression profile. Cluster Analysis of Gene Expression Dynamics (CAGED), a clustering technique proposed by *Ramoni et al*.^20^, uses Bayesian method to model gene-expression dynamics using auto-regressive equations. TimeClust^21^ uses temporal gene expression profiles to produce clusters. TMixClust^22^, Dirichlet process Gaussian process mixture model^14^ are some of the significant clustering approaches containing methods with non-parametric models.

The analysis of time-series gene expression modules has helped us unravel major biological complications. It has given us serious insights into disease progression^23^, biomarker discovery^24^, identification of hub genes^25^, cell cycle progression^26^, cancer classification^27^ and several other bio-medically important processes. Moreover, the advancements in information system infrastructure have facilitated creating time-series schemes more feasible method for studying complex psychological phenomena. Multiple tools are available to provide us with an enriched network and pathway analysis of these gene modules. Using such tools enables us for further analysis and deeper observation of the biological mechanisms.

In this article, we have proposed a framework to discover key transcriptomic regulators and key modules containing majority of the key regulators from time-series microarray gene expression data for pancreatic ductal adenocarcinoma. Initially, differentially expressed genes (DEGs) were identified by analyzing empirical Bayes statistics on multivariate time-course gene expression data of pancreatic cancer in R (splineTimeR)^28^. Also top 100 DEGs at each time point were analyzed using an R/Bioconductor software package called Linear Models for Microarray Data (Limma)^29^. Clustering on the DEGs to form gene modules based on similar responses was achieved by a non-parametric model, Dirichlet process Gaussian process mixture model^14^ in Python. A web tool, REGulator-Gene Association Enrichment (REGGAE)^30^ was used for the identification of key transcriptional regulators along with the number of targets for each of them from the list of DEGs.

Most of the experimental gene expression studies only focus on the determination of DEGs by handling them as independent events and do not investigate the co-action of identified genes. Reconstruction of the possible gene association network (GAN) among DEGs helps us find genes that take part in the interaction network of studied phenotype. Therefore, reconstruction of GAN followed by the identification of top nodes in the network was also performed. Prediction of gene functions of the top genes in the interaction network has also been carried out using GeneMANIA prediction server^31^. We have identified the key gene modules from the list of modules obtained from the clustering. Subsequently, transcriptomic regulatory genes were also detected against a curated database of DNA-binding RNA polymerase II TF (DbTF) using TFcheckpoint^32^. Furthermore, biological significance like Kyoto Encyclopedia of Genes and Genomes (KEGG) pathway, Gene Ontology (GO) and gene-disease associations of the gene modules were also observed using a gene set enrichment analysis tool, Enrichr^33^.

## 1 Results and Discussion

This section gives us insight about the detailed findings of our present work.

### 1.1 Evaluation of Differential Expression

We have processed 42412 genes with their normalized expression values from methods discussed in section 2.1. These genes along with their normalized expression values were analyzed for differential expression. Differential gene expression analysis on genes has been executed using splineTimeR as described in section 2.2. We obtained 1397 genes that were differentially expressed by using adjusted *P*-value ≤ 0.05 using the correction method of Benjamini-Hochberg (BH)^34^ with 4 as the optimum degree of freedom. Top 20 genes (DEG’s) from the pancreatic ductal adenocarcinoma dataset with significant expression value changes were ‘Hs.7413’, ‘CYP26B1’, ‘NPPB’, ‘SFRP4’, ‘Hs.562219’, ‘C12ORF46’, ‘SMAD3’, ‘RN7SK’, ‘RARRES1’, ‘ITGA4’, ‘CLSTN2’, ‘DKK1’, ‘CYP26A1’, ‘EPDR1’, ‘RARB’, ‘LOC340598’, ‘TMEM16C’, ‘INMT’, ‘ACTC1’, ‘C21ORF7’. Figure 8 represents the box and violin plot of the cubic spline normalized gene expressions of samples for each condition at individual time point of the differentially expressed genes. We have also performed DE analysis for all the genes at each time point to distinguish the genes expressed at each time point using Limma package in R. Table 4 shows the top 5 DEGs at each time point, including the top 5 genes of the final DEG list, found using Limma.

We have analyzed the genes in Limma by applying empirical Bayes smoothing on the estimated fold-changes and standard errors from a linear model fitting. We have taken the top 100 genes from each time point using BH adjustment on adj. *P*-values to determine DEGs. The common differentially expressed genes at each time point have been identified. DEG list of the top 100 genes had the highest similarity of 59% with the top 100 genes of the final list at 168 hours amongst all the time points, followed by 41% at 24 hours. Figure 1 represents the Venn diagram of the top 100 DEGs at each time point with no genes at the intersection of all time points and DEG_*t* stands for the top 100 DEGs at time point *t*, where *t* ∈ {30 Minutes, 4 Hours, 12 Hours, 24 Hours, 168 Hours}. DEGs from splineTimeR and top 1397 genes from the final table of Limma were 100% similar.

**Figure 1.**
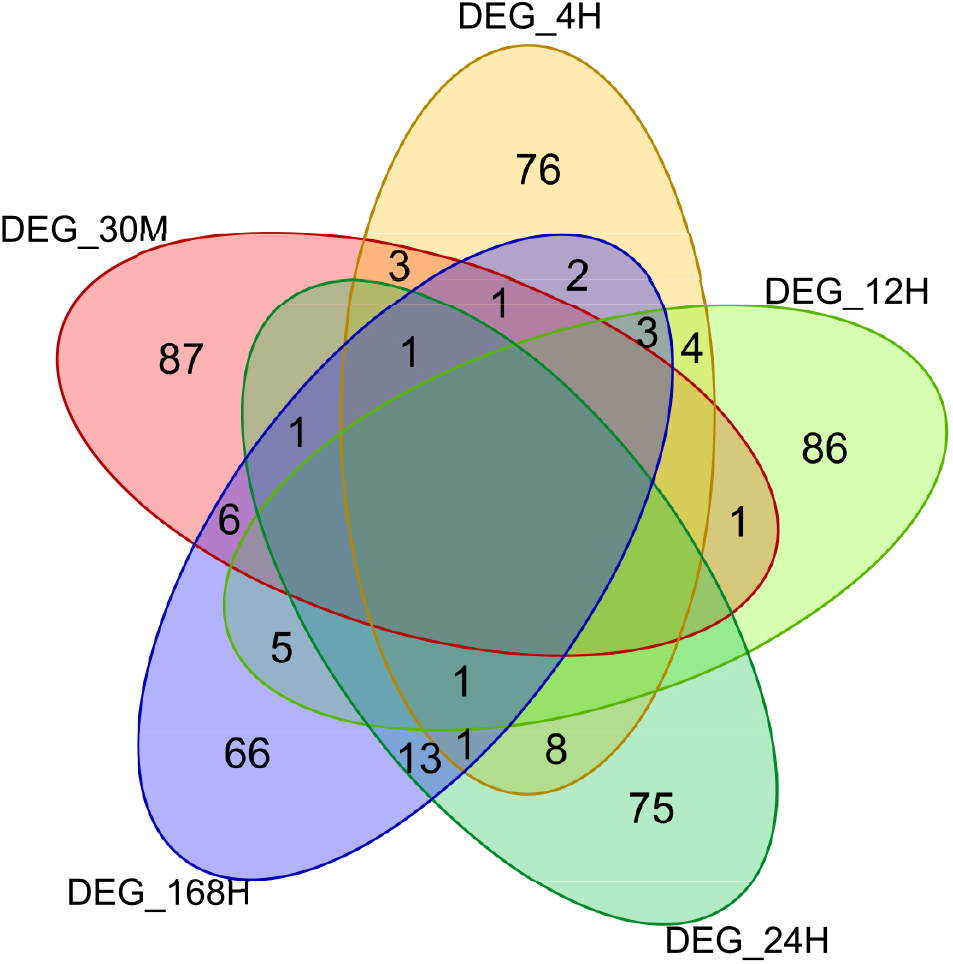
Venn Diagram of number of DEG’s at each time point.

### 1.2 Gene Network Reconstruction and Gene Function Prediction

Adopting the methodologies discussed in section 2.3, we have reconstructed the gene association network (GAN). We have obtained two GAN’s. We got 45550 significant edges with 1107 nodes for cutoff probability 0.8 and 34080 significant edges with 1048 nodes for cutoff probability 0.9, which corresponded to 4.67% and 3.5% of possible edges respectively. Figure 2 depicts the reconstructed GAN with a probability cutoff set to 0.9 for the top 150 selected edges based on a higher partial correlation score. Genes with higher degrees in the whole reconstructed GAN are displayed in dark color. From this figure, it can be observed that ‘CUEDC1’, ‘ABCA6’, ‘MRPL50’, ‘LYPD3’, ‘KRT19’, ‘OLFML3’, ‘LGALS3’, ‘Hs.540914’, ‘LOC285074’, ‘TBPL1’ are the top 10 genes having extremely high connection with others within the GAN. We have also generated reconstructed GAN highlighting the betweenness and closeness centralities for the top 150 selected edges based on higher partial correlation scores which are shown in **Supplementary File 1**. It was discovered that ‘CUEDC1’, ‘ABCA6’, ‘MRPL50’, ‘LYPD3’, ‘KRT19’, ‘OLFML3’, ‘TBPL1’ were the seven common genes which ranks within top 10 using all of these three centrality measures.

**Figure 2.**
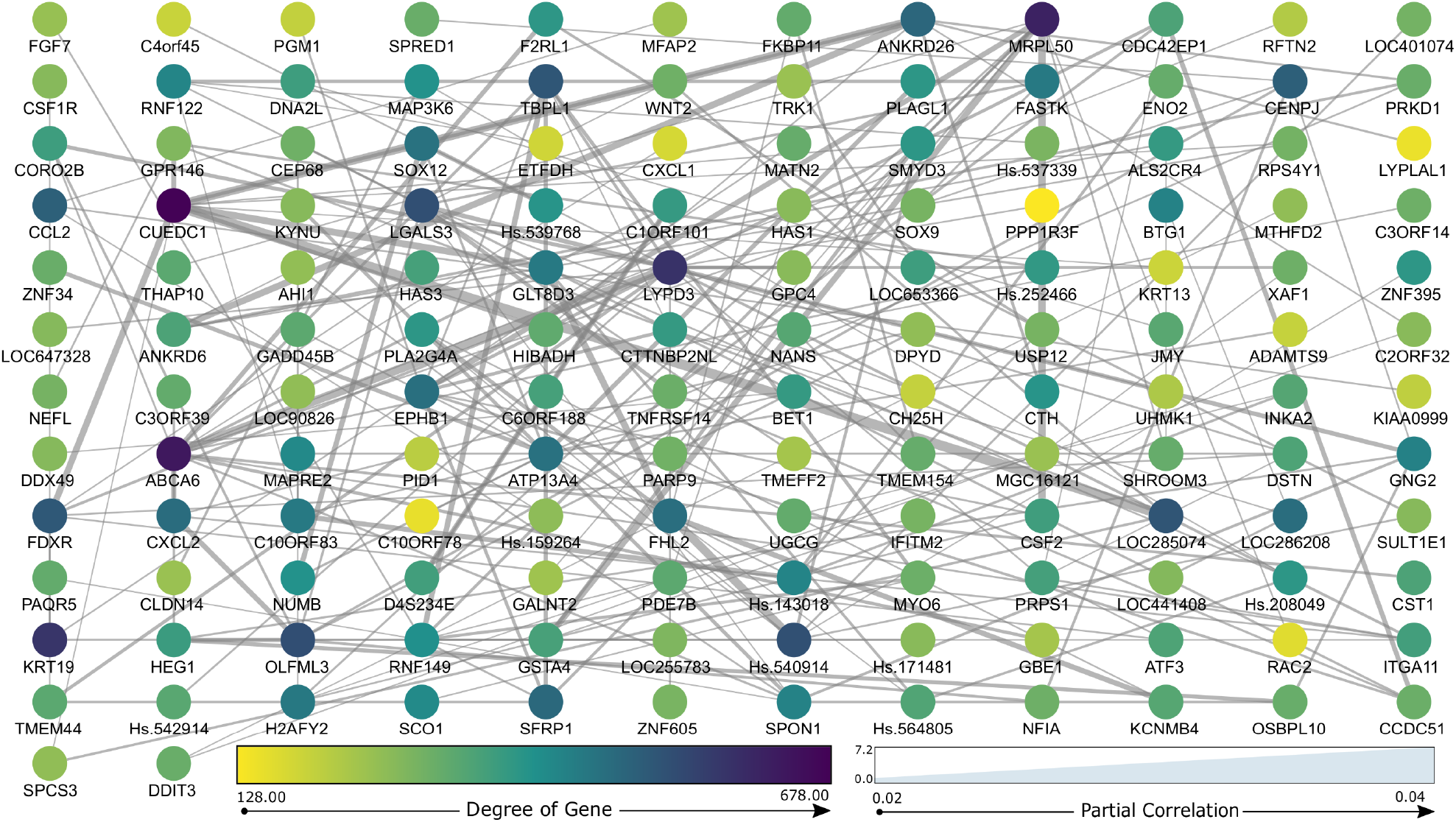
Reconstructed Gene Association Network using splineNetRecon with top 150 edges based on partial correlation score.

We have also extracted the top 150 genes from the reconstructed GAN with 0.9 as the probability cutoff for prediction of their gene function, chosen based on the aggregation of three centrality measures: degree, betweenness, and closeness centrality. The top 15 genes among them include ‘CUEDC1’, ‘ABCA6’, ‘MRPL50’, ‘LYPD3’, ‘KRT19’, ‘OLFML3’, ‘LGALS3’, ‘TBPL1’, ‘LOC285074’, ‘ANKRD26’, ‘FDXR’, ‘Hs.540914’, ‘CCL2’, ‘CENPJ’, and ‘CXCL2’. We utilized the list of genes to obtain their gene function using GeneMANIA web server in Cytoscape app. GeneMANIA resulted in the creation of a weighted connected network of query genes including the suggested genes by the algorithm. The resultant network is demonstrated in Figure 3. We have also obtained the predicted gene functions by GeneMANIA. Table 1 provides us with comprehensive details of the top 5 functions of the resultant genes.

**Table 1.** GeneMANIA predicted functions.

|  | Function | FDR | Coverage |
| --- | --- | --- | --- |
| 1. | Cellular response to tumor necrosis factor | 0.0015 | 8/74 |
| 2. | Response to tumor necrosis factor | 0.0027 | 8/87 |
| 3. | Response to transforming growth factor beta | 0.025 | 9/169 |
| 4. | Cellular response to transforming growth factor beta stimulus | 0.025 | 9/169 |
| 5. | Cellular response to interleukin-1 | 0.026 | 5/36 |

**Figure 3.**
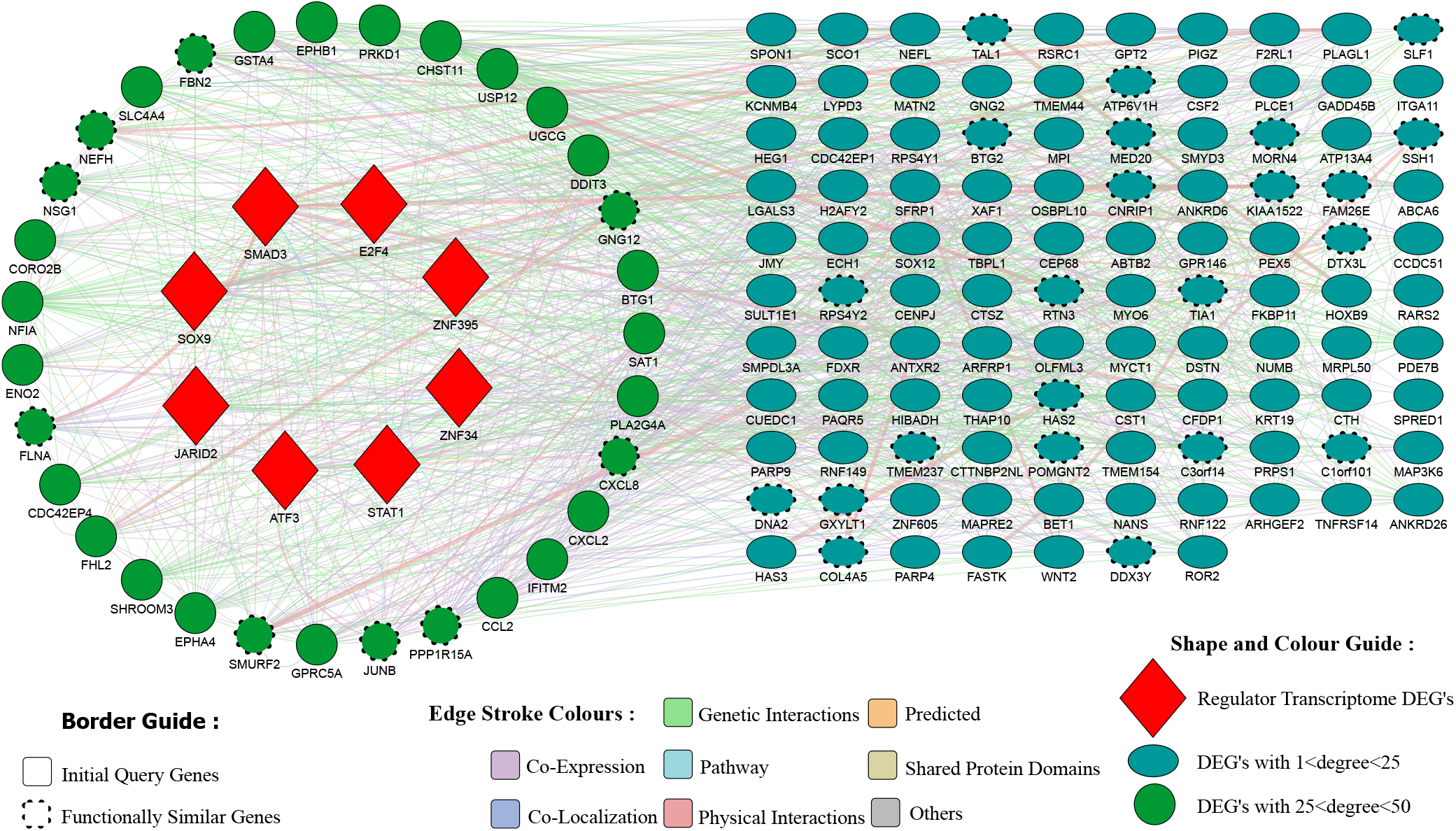
Gene network created from GeneMANIA web server.

In table 1, the coverage field defines a ratio of the number of genes in the resultant network to that in the genome, and FDR represents the False Discovery Rate generated from GeneMANIA algorithm. The top 50 genes have been chosen from the resultant ranked gene list provided by GeneMANIA with the assigned score for finding their gene-disease associations. It has been observed that ‘CSF2’, ‘HOXB9’, ‘ITGA11’, ‘NUMB’ genes are associated with Pancreatic Ductal Adenocarcinoma according to DisGeNET^35^ web server. The resultant gene list also contained 8 DNA-binding TF’s including ‘SMAD3’, ‘STAT1’, ‘ZNF34’, ‘ATF3’, ‘ZNF395’, ‘JARID2’, ‘E2F4’, ‘SOX9’. Results are described in section 1.3.

### 1.3 Transcriptional Regulators and DNA-binding Transcription Factor Identification

We have identified key transcriptional regulators from the generated DEG list using REGGAE^30^. We obtained a ranked list of 66 key transcriptional regulators from an RTI collection which contains a list of regulators for each deregulated gene that may influence its expression. Genes were ranked according to the score provided by the REGGAE algorithm. The top 5 genes from the list of key TR’s are ‘GATA6’, ‘NFYB’, ‘IRF1’, ‘TRIM22’, ‘SREBF1’. Among the generated list of key TR’s, 5 TR’s were not from the genes of the DEG list implying multiple regulators influencing a deregulated gene. Those 5 TR’s are ‘PEX2’, ‘FOXO1’, ‘RREB1’, ‘HEXIM1’, ‘ZNF280D’.

Additionally, we have treated the genes in the key TR list with a curated database of specific DbTFs: TFcheckpoint^32^, to find proteins playing a central role in determining which genes were transcribed. We retrieved 47 DNA-binding RNA polymerase II TFs among 66 key TR’s. This 47 DbTFs contained 10 proteins that were directly associated with Pancreatic Ductal Adenocarcinoma (PDAC) disease. The proteins, according to DisGeNET, associated with this disease are ‘FOXO1’, ‘SOX9’, ‘GATA6’, ‘SMAD3’, ‘NFKB1’, ‘KLF6’, ‘TBX3’, ‘SREBF1’, ‘NR4A2’, ‘TCF3’. In table 1, the coverage field defines a ratio of the number of genes in the resultant network to that in the genome, and FDR represents the False Discovery Rate generated from the GeneMANIA algorithm. The top 50 genes were chosen from the resultant ranked gene list provided by GeneMANIA with the assigned score for finding their gene-disease associations. It has been observed that ‘CSF2’, ‘HOXB9’, ‘ITGA11’, ‘NUMB’ genes are associated with Pancreatic Ductal Adenocarcinoma according to DisGeNET^35^ web server. The resultant gene list also contained 8 DNA-binding TF’s including ‘SMAD3’, ‘STAT1’, ‘ZNF34’, ‘ATF3’, ‘ZNF395’, ‘JARID2’, ‘E2F4’, ‘SOX9’. Results are described in section 1.3.

### 1.4 Gene Modules Identification and Determination of Key Gene Modules

After discovering DEGs and DbTFs from the list of DEGs, we have applied the Dirichlet Process Gaussian Process mixture model clustering to obtain robust and accurate gene modules incorporating their expression values at different time points. The optimal number of clusters obtained by using default conditions as discussed in section 2.4 was 32, with 136 and 10 genes being the highest and the lowest number of genes in a module among all the modules, respectively. Figure 4 shows the complex heatmap denoting the posterior distribution of the probability that two genes are being co-clustered.

**Figure 4.**
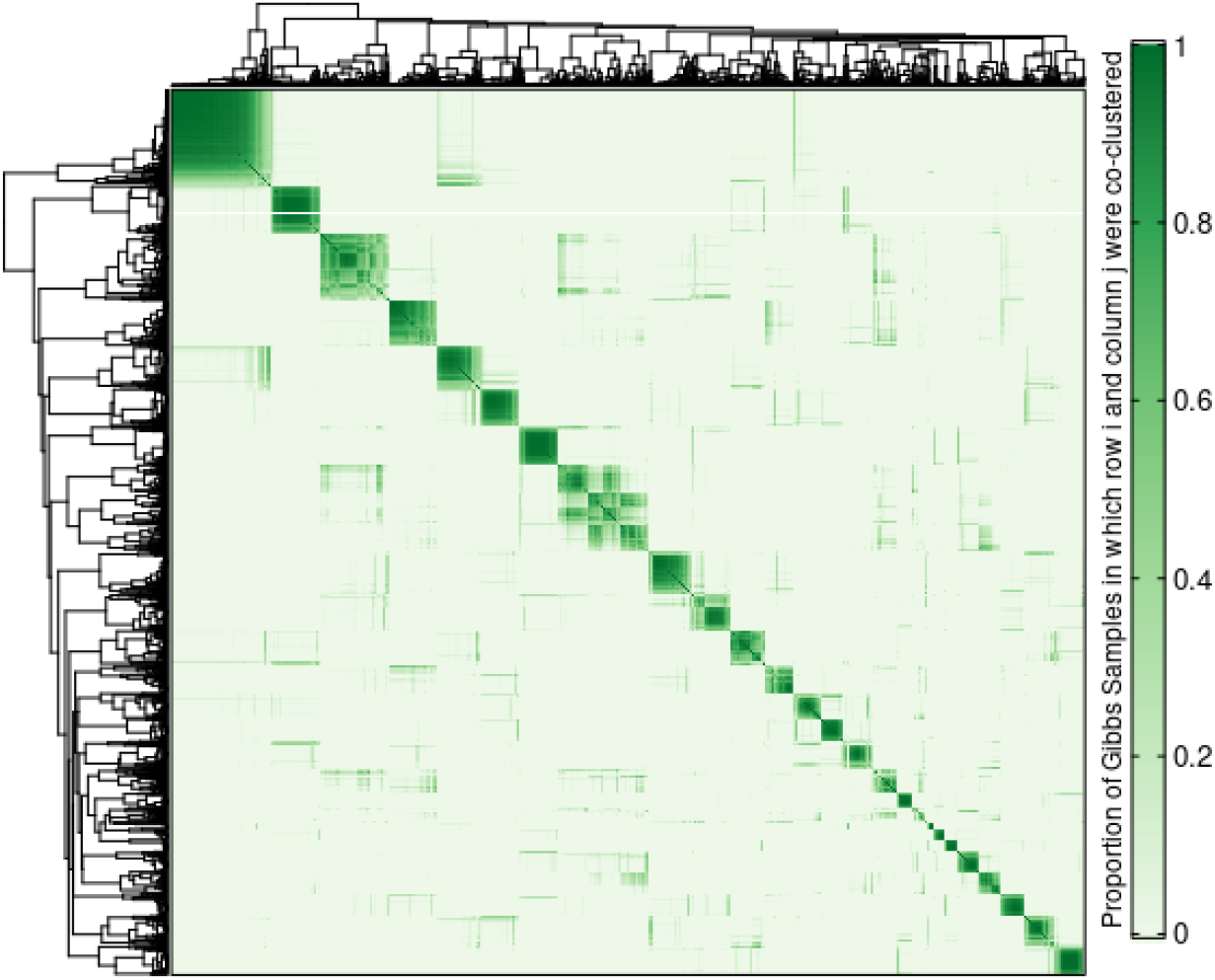
Heatmap of posterior similarity matrix from clustering.

Subsequently, we have used the modules and the genes inside each module to find the key modules. We have assigned rank to the clusters according to the descending order of the number of regulator genes inside each gene module. The top 4 modules with the highest number of regulators have been considered as the key modules. We have discovered that ‘gene module’ 2 contains 7 key TRs (now onward will be referred to as *‘Key Cluster 1’*), ‘gene module’ 1 contains 6 key TRs (now onward will be referred to as *‘Key Cluster 2’*), ‘gene module’ 9 contains 6 key TRs (now onward will be referred to as *‘Key Cluster 3’*) and ‘gene module’ 26 contains 6 key TRs (now onward will be referred to as *‘Key Cluster 4’*) are the 4 key modules. The key modules had 78, 136, 50, 64 number of genes in total for each cluster, respectively. Here, we also found out that only 24 gene modules out of 32 modules contained all the key TRs.

Furthermore, we have studied the normalized gene expressions in each of the 4 key modules for 5 time points to understand the magnitude of variation of expression values in the modules. Figure 5 gives us a visual representation of the average fold-change in gene expression along with individual gene trajectory for the 4 key clusters.

**Figure 5.**
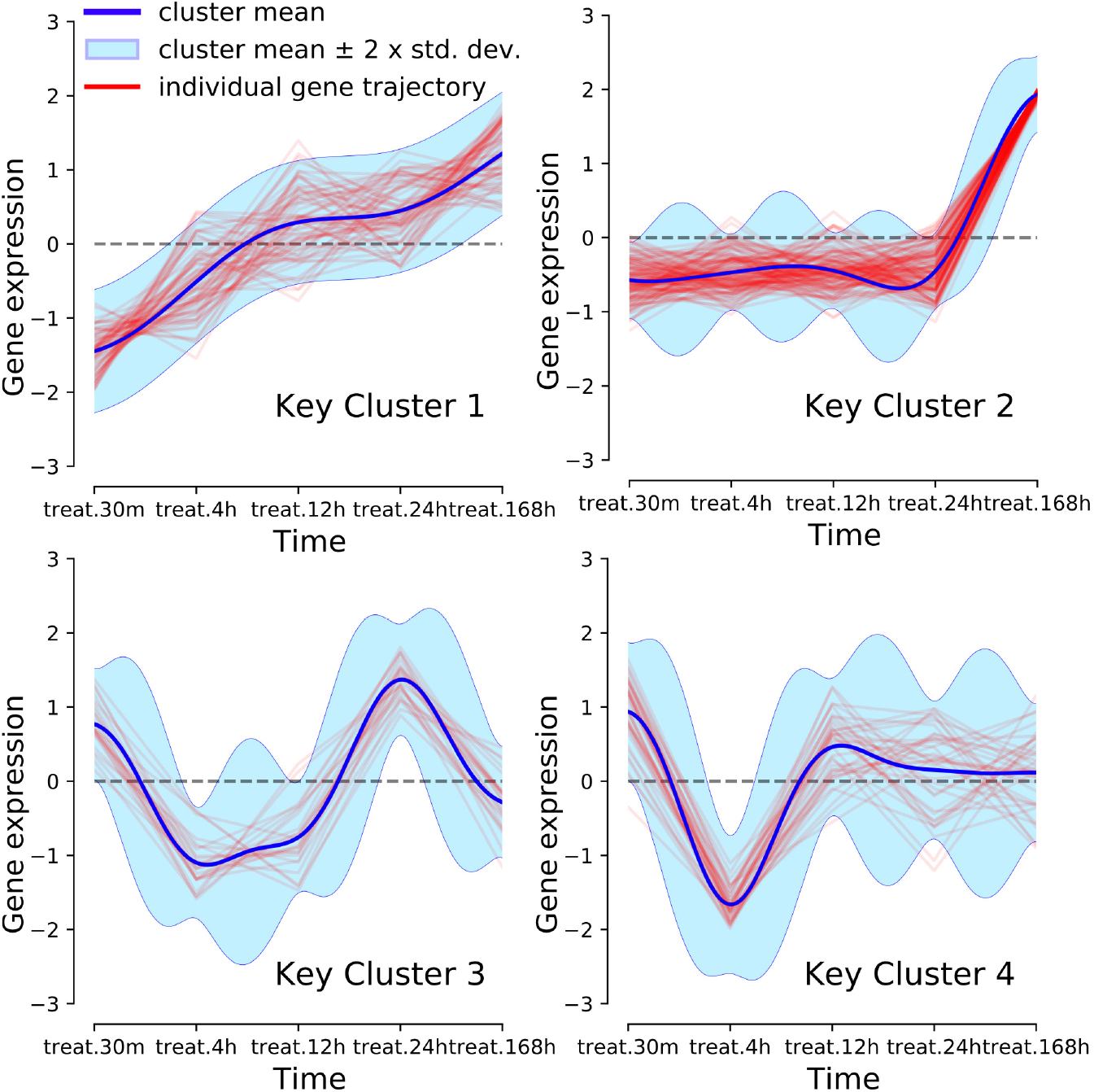
Statistics of gene expression in top 4 key modules (Key Clusters).

### 1.5 Biological Significance Analysis

We have performed several analysis on the key modules in order to gain more insights about the pathways and biological processes (BP) of the involved genes inside the key modules. Additionally, we have identified key TRs in each of the key module. We have identified KEGG pathways, biological processes (BP) details of the Gene Ontology terms and disease-gene associations using DisGeNet database of Enrichr web server^33^. These information along with all the regulator genes and DbTF among the key TRs have been summarized in Table 2, where genes in bold face have been used to represent DbTFs among the TRs. Figure 6 shows the top ranked (based on *p*-value) KEGG pathways (6 A), biological processes (6 B) and drugs/disease association (6 C) enrichment of genes in key cluster 1. These analysis for the other 3 key modules has been attached in the **Supplementary File 1**.

**Table 2.**
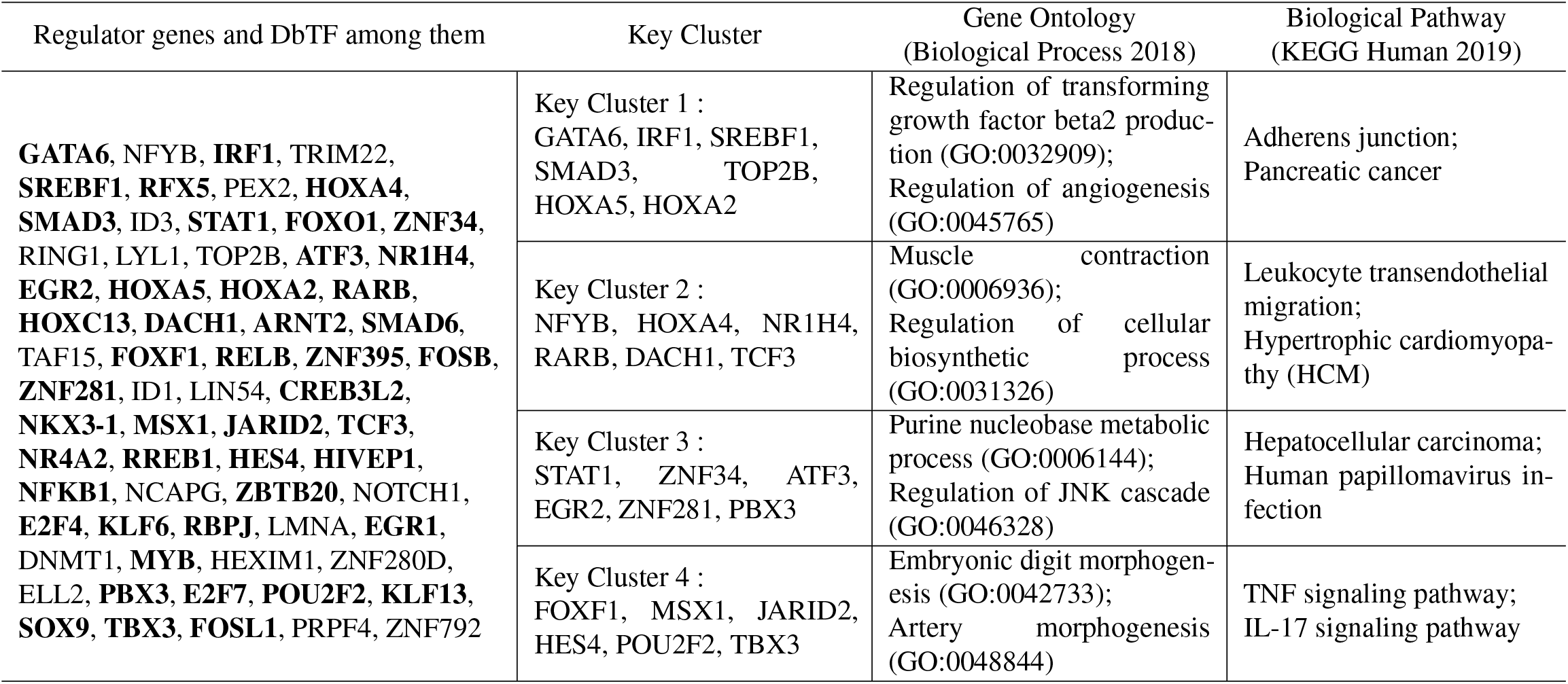
Biological significance analysis of key regulators in each key module.

**Figure 6.**
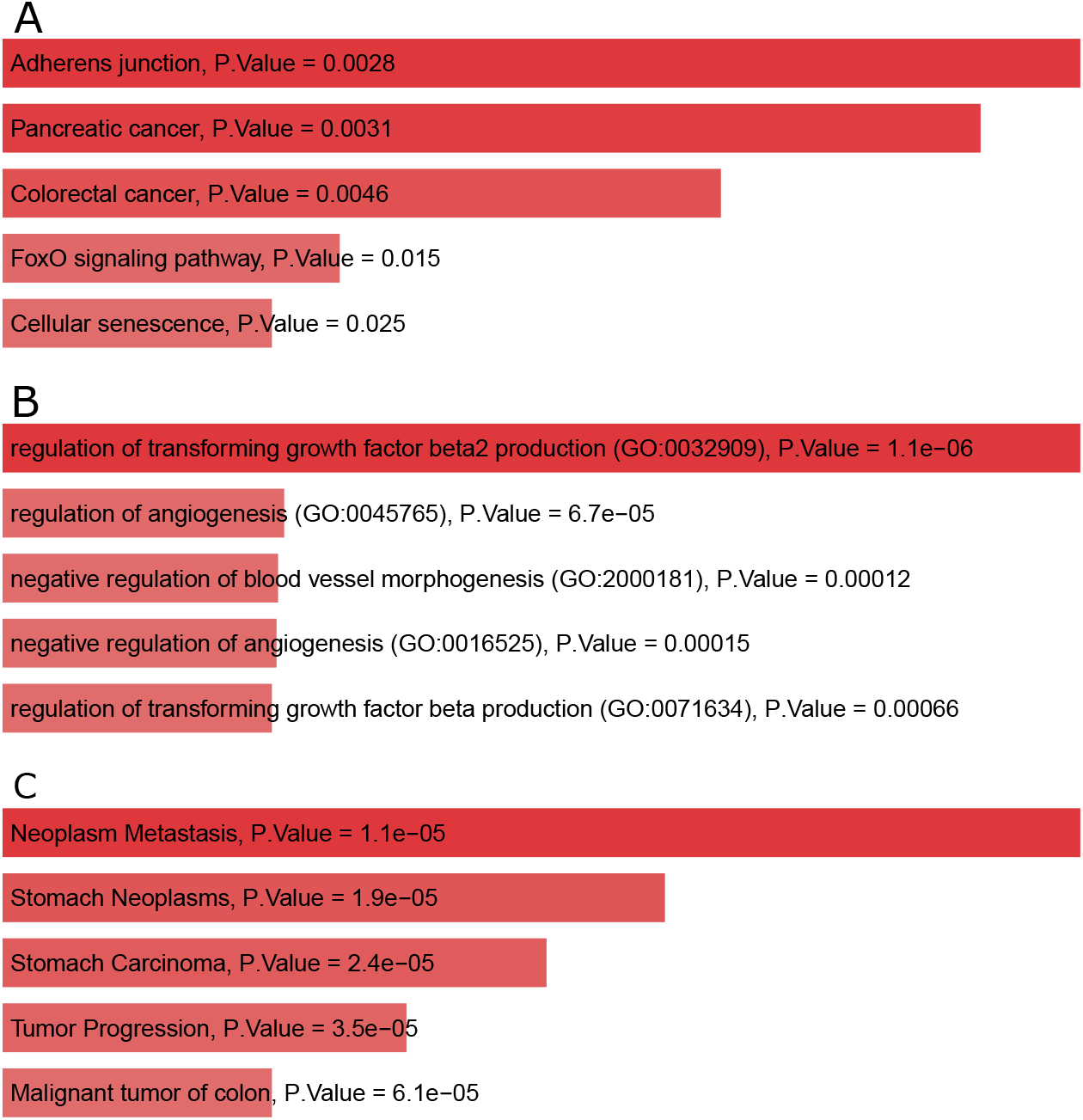
A. KEGG Pathway of *Key Module 1*, B. Biological Process of *Key Module 1*, C. DisGeNet analysis of *Key Module 1*.

Moreover, we have observed the top 5 KEGG pathways and GO terms (BP) of each of the key modules to understand their relevance and significance in PDAC. We have found several important factors essential for PDAC disease among the observed results. We observed that key module 1 regulates Transforming Growth Factor (TGF) beta2 production which leads to high damage for over-expressing TGF-*β* in PDAC^36^. In key module 2, ‘MYL7’, ‘SHC3’, ‘ITGA4’, ‘COL4A4’, ‘ITGA1’, ‘FLNA’, ‘MYL9’ genes are significant in focal adhesion pathway which plays a key role being over-expressed and highly active in PDAC^37^. In a publication, Gordon et. al. demonstrated that several bone morphogenesis signaling component targets has been up-regulated in pancreatic cancer tissues including genes like ‘SFRP4’, ‘CYP26B1’, ‘RARB’^38^. Key module 3 is responsible for the regulation of c-Jun N-terminal kinase cascade regulation for the presence of ‘DUSP10’, ‘GADD45A’, ‘ANKRD6’ genes which is one of the pathways activated in PDAC^39^. Regulation of the p38MAPK cascade pathway in key module 3 is involved in Interleukin 1 alpha (IL-1*α*) expression by PDAC^40^. We found that Interleukin-17 signaling pathway promoted changes from chronic pancreatitis to pancreatic cancer accompanied by ‘CSF2’, ‘CCL20’, ‘TNFAIP3’ genes in key module 4^41^. Importance of tumor necrosis factor, notch signaling pathway are also significant for promoting PDAC^42,43^. Finally, we found that ‘TGFB2’, ‘SMAD3’, ‘RAC2’ genes in key module 1 were responsible for pancreatic cancer pathways.

Figure 7 shows the top 15 gene ontology terms (biological processes) performed by the genes in the ‘key module 1’ with their proportion of counts in the module. These GO terms have been selected according to the lowest *p*-values. This analysis for the other 3 key modules has been attached in the **Supplementary File 1**.

**Figure 7.**
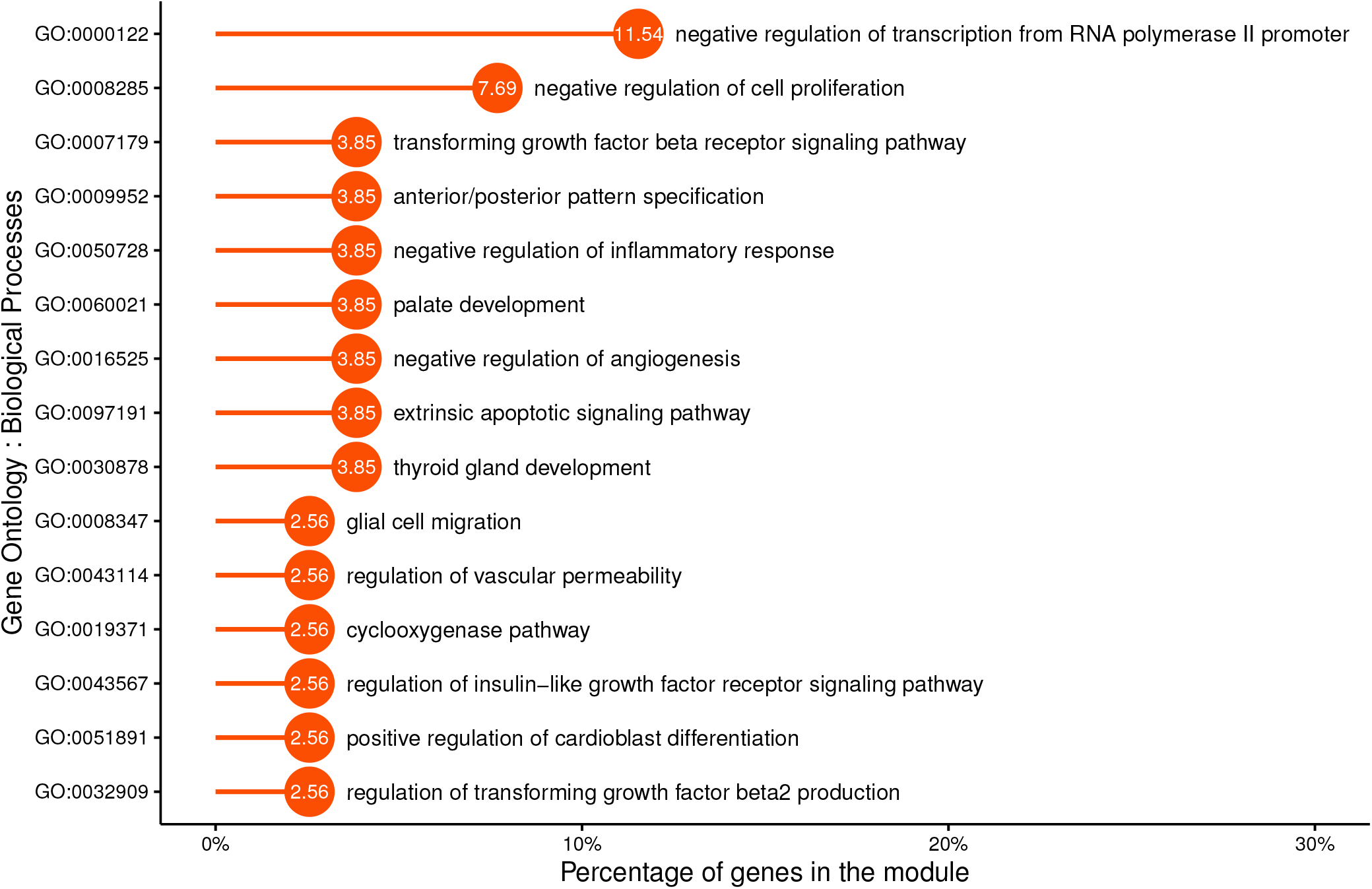
Lollipop plot describing percentage of gene in *key module 1* contributing to top 15 Gene Ontology terms.

**Figure 8.**
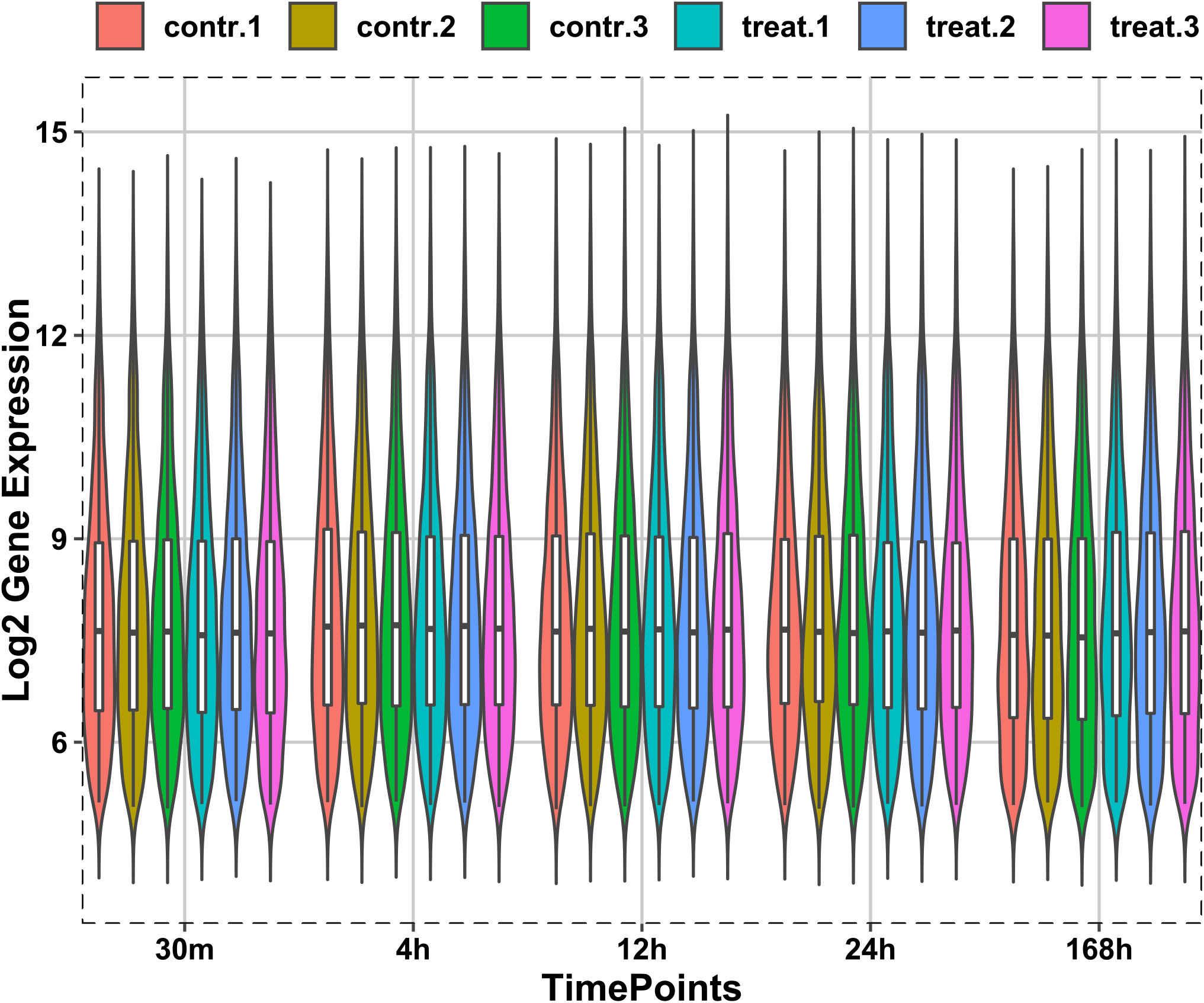
Box and Violin plots of cubic spline normalized gene expression of the differentially expressed genes.

Additionally, we have analyzed genes in the top 4 key modules for their direct association with PDAC. We have observed the gene list from each key module against DisGeNet web server. We have found that 6, 8, 3, 7 genes were directly associated with PDAC in the key modules 1, 2, 3, 4, respectively. The names of the genes have been tabulated in Table 3.

**Table 3.** Genes related to PDAC in each key module.

| Key Module 1 | Key Module 2 | Key Module 3 | Key Module 4 |
| --- | --- | --- | --- |
| GATA6,<br>IGFBP3,<br>SMAD3,<br>SREBF1,<br>TIMP3,<br>GPRC5A | IGF2BP3,<br>CNN2, IDH1,<br>CXCL12,<br>MMP11,<br>SFRP4, TCF3,<br>NCOA3 | ROR2,<br>CLIC3,<br>ITGA11 | LGALS3,<br>CSF2, ISG20,<br>TBX3,<br>NUMB,<br>MTSS1,<br>KHDRBS1 |

**Table 4.** Differentially Expressed Genes using Limma.

| 30 Minutes | 4 Hours | 12 Hours | 24 Hours | 168 Hours | Final DEG's |
| --- | --- | --- | --- | --- | --- |
| SNAR-A1 | CXCL2 | FKBP11 | CYP26B1 | CYP26B1 | Hs.7413 |
| HMOX1 | CCL20 | MYH10 | SYNPO2L | FHOD3 | CYP26B1 |
| LFNG | DHRS3 | CRLF1 | CXCL1 | NPPB | NPPB |
| BCYRN1 | IL8 | SLC12A8 | NPPB | ACTC1 | SFRP4 |
| LOC255783 | SMAD3 | Hs.559673 | LOC340598 | SFRP4 | Hs.562219 |

## 2 Methods

The present section provides the overview of our systematic approach on data collection, data preprocessing and the overall framework for the different methodologies used in our present analysis.

### 2.1 Data Preparation

Gene Expression Omnibus (GEO)^44^ has been used to collect Pancreatic ductal adenocarcinoma microarray dataset (GSE14426), which contains treated pancreatic stellate cell line raw counts. The dataset has 48701 genes and 30 samples of each gene along with their periodic gene expression changes. Samples include cubic spline-normalized intensity values of two conditions, viz., pancreatic stellate cell line before and after being treated with 1-molecular concentration of all-trans retinoic acid (ATRA) on plastic. The gene expression values were recorded at 5 time points (30 min, 4 hours, 12 hours, 24 hours and 168 hours) with 3 replicates at each time point. All expression values were *log*_2_-transformed for better performance in finding DEGs.

The R package *org*.*Hs*.*eg*.*db*, genome-wide annotation database for Homo Sapiens^45^ was used to map the Illumina gene identifiers from the raw data file to their official gene symbols along with the Illumina human platform information (GPL6102) from GEO. We have obtained the final set of genes after mapping with official gene symbols, considering all gene names found from 1:1 as well as 1:many mapping and aggregating expression values of duplicate genes from the total number of genes.

### 2.2 Differential Expression Analysis

In order to analyze and extract the genes with significant gene expression changes, differential gene expression analysis has been performed using the R/Bioconductor *splineTimeR* package. SplineTimeR operates on the values obtained from the parameters of a fitted natural cubic spline regression model. The regression model is fitted with time-dependent gene expression data for two expression groups (e.g. control and treated)^28^. The differential expression of a gene has been discovered by applying empirical Bayes moderate F-statistics on the coefficient differences of the spline regression model. The mathematical natural cubic spline regression model is defined as:

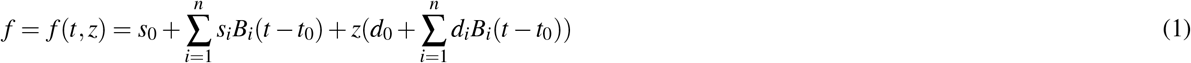

where *s*_*i*_ and *d*_*i*_ is the reference group’s spline coefficient and compared group’s differential spline coefficient, respectively and *B*_*i*_(*t* − *t*_0_) is spline base function ∀_*i*_ = 1, 2, …, *n* with *t*_0_ being the first measurement time.

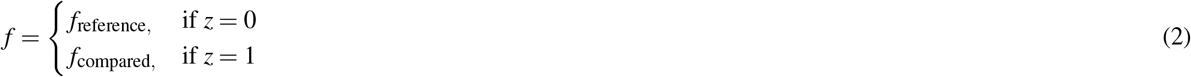

The detection of DEGs using splineTimeR has been carried out by setting the Benjamini-Hochberg^34^ adjusted *P*-value threshold to 0.05 and with a degree of freedom as 4 for all genes. Top regulated genes have been identified as differentially expressed and used for further analysis. Figure 8 represents the box and violin plot of the cubic spline normalized gene expressions of samples for each condition at the individual time point of the differentially expressed genes.

Additionally, the Limma^29^ software package has been utilized to find top DEG’s at each time point considering the replicates of that particular time point. Limma fits linear models to gene expression value matrix to assess differential expression by taking care of the differences among gene sample conditions. A linear model can be defined as:

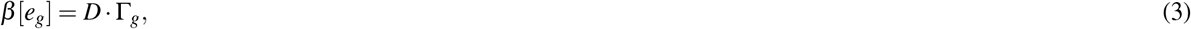

where *e*_*g*_ represents a vector of expression values and *D* represents the design matrix which maps these vector to some coefficient of interest Γ_*g*_ for each gene *g*. A contrast matrix was also used for computing *log*_2_-fold-changes and *t*-statistics by allowing all possible pairwise comparisons between gene samples. Furthermore, test-statistics are obtained to perform gene ranking. Normalized expression values for 3 replicates at each of the 5 time points were used for differential expression analysis.

Table 4 shows top 5 DEGs at each time point, including the top 5 genes of the final DEG list, found using Limma.

### 2.3 Gene Association Network Reconstruction and Prediction of Gene Function

In this work, we have performed a gene association network (GAN) reconstruction from time-course data of the prior selected DEGs using a regularized dynamic partial correlation method. GeneNet^46^ R-package has been used to analyze covariance matrices with selected shrinkage method as dynamic. Analyses have been performed with a posterior probability of 0.8, and 0.9. GAN reconstruction has been implemented using splineNetRecon function from the splineTimeR package to identify regulatory association between genes under a specific condition (Treatment)^28^. Top significant edges have been identified based on their cutoff posterior probability.

Important genes have been identified by analyzing the overall graph topology in the resultant GAN using the commonly used centrality measures: degree, betweenness and closeness centrality. These centrality measures were combined for selecting the top 150 ranked genes by taking the average of the quantile normalized values of these measures to perform gene function prediction.

GeneMANIA^31^ provides us with a very adjustable interface to obtain possible connections between the query genes by searching several prominent networks containing gene function, proteomic data. GeneMANIA operates on a ridge regression derived fast heuristic algorithm to integrate multiple functional gene association networks using label propagation. GeneMANIA weights data sources based on the connectivity strength of all genes with each other in the query list and suggests relatively similar non-queried genes and their connection types. It returns an interactive functional gene association network of all resultant genes and their relationship. It also returns functions of interest based on non-negative weights of gene sources (i.e. association of two genes) by considering the problem of prediction of gene function as a binary classification problem. GeneMANIA further returns a ranked gene list presumably sharing phenotypes with query list genes based on it’s large and diverse data. All 150 genes from the ranked list of GAN were analyzed for gene function using the GeneMANIA Cytoscape app.

### 2.4 Finding Gene Modules from Time-Series data

Piece-wise Aggregate Approximation (PAA) has been used from the *TSrepr*^47^ package in R for dimensionality reduction of multivariate time-series data. The DEGs obtained from the dataset contained 3 replicate values at each time point, which have been converted into univariate expression values for each of the 5 time points using the mean function of PAA. These expression values were clustered to form gene modules.

We have used a model-based approach, the Dirichlet Process Gaussian Process mixture model (DPGP), presented by McDowell *et al*. to find gene modules from univariate time-course expression data^14^. It simultaneously models cluster number with a Dirichlet process and also uses temporal dependencies as a Gaussian Process. DPGP uses a Bayesian non-parametric model for time course paths *P* ∈ ℝ^*N*×*T*^, where *N* is the number of genes and *T* is the number of time points. A generative Dirichlet process (DP) mixture model is defined as

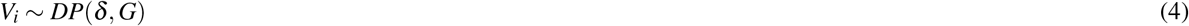

Here, *DP* represents a draw which generates latent variables *V*_*i*_ for a cluster *i* with *δ* being the concentration parameter and *G* being the base distribution, and the observation distribution *M*_*j*_ ∼*p*(.| *V*_*i*_) is specified with a Gaussian Process (GP) for gene *j* ∈ {1, 2, …., *N*}. Cluster-specific parameter values *V*_*i*_ is obtained from conditional probability *p*(*V*_*i*_ |*V*_¬ *i*_) on all other variables except *V*_*i*_ by integrating draws from *DP*. The within-cluster parameters of the trajectories for a cluster *i*, i.e. 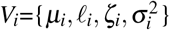 is described as *µ* _*i*_ ∼ *GP*(*µ*_*0*_, *L*) with *l* _*i*_, *ζ* _*i*_ ∼ ln 𝒩 (0, 1) and 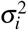 being an inverse gamma function of shape and rate of the trajectory. DPGP uses a cluster-specific positive definite Gram matrix *L* quantifying similarity between each and every time point *T*_*x*_, *T*_*x′*_ for a cluster *i* which is defined as

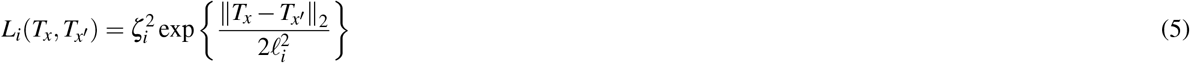

The hyperparameter *l* _*i*_, *ζ*_*i*_ corresponds to input space distance among data points having correlated output (*characteristic length scale*), gene expression path variance over time (*signal variance*), respectively. Thus, the probability distribution of each observation *M*_*j*_ corresponding GP for a specific cluster including unique marginal variance is characterized by

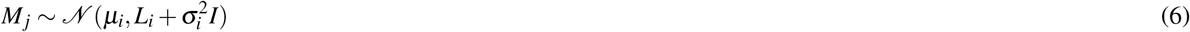

DPGP uses Markov Chain Monte Carlo Method (MCMC)^48^ to assess the posterior distribution of the parameters used in the model. In MCMC, “Algorithm 8” of Neal’s Gibbs Sampling^49^ has been implemented to estimate the posterior distribution of the probability that a cluster *i* containing trajectory of a gene *j* corresponding to DP. We ran the MCMC for 1000 iterations which is also the default number of iterations, among which the first 48% are divided into two burn-in phases equally. Initially, every gene is assigned to the clusters of their own. In burn-in phases, a cluster is chosen at each iteration for a gene based on its likelihood to fit a GP having cluster mean and covariance as initial parameters. After the second burn-in phase posterior probability of the kernel hyperparameters was computed. MCMC generates a state sequence from Gibbs’ sampler in which each state contains a section of genes into each cluster. Maximum a *posteriori* clustering or partition with the maximum value of posterior probability has been chosen to summarize results. We have used a Python implementation of DPGP from the command line interface and resultant clusters of genes have been used for further analysis.

### 2.5 Key module Identification and Biological Significance Analysis

This subsection provides information about approaches that have been used to find biological significance of our obtained results.

#### 2.5.1 Finding Transcriptional Regulators

We have analyzed DEGs along with their expression values between two sample points to identify key transcriptional regulators (TR) using REGGAE^30^. It operates by using Kolmogorov-Smirnov-like test statistics along with an implicit combination of Regulator Target Interactions (RTI’s) for the prioritizing influence of TR’s. It returns a ranked list of key TRs with p-value aggregations.

#### 2.5.2 DNA-binding Transcription Factor Identification

Key TRs have been observed against a cumulative and high-quality knowledge source of genome-scale information referring their biological function potentiality as DNA Binding Transcription Factors (DbTF) using TFcheckpoint^32^.

#### 2.5.3 Identifying Key Modules

Gene modules have been studied and ranked for having the highest number of key TRs. Gene modules having the highest number of key TRs were identified as the key modules. Further analysis of the top 3 modules was carried out.

#### 2.5.4 Gene Set Enrichment Analysis

Top 3 key modules have been analyzed to discover biological processes and pathway analysis for the involved genes inside each module. Biological aspects like gene ontology, disease-gene associations and pathway analysis were determined using Enrichr^33^.

## 3 Conclusion

In this article, we have designed a framework for analyzing top regulated genes from multivariate time-series Pancreatic Ductal Adenocarcinoma (PDAC) microarray data to identify key gene modules, DNA binding Transcription Factors (DbTF) among key Transcriptomic Regulators (TR), and gene function prediction of potential crucial genes in construction of Gene Association Networks (GAN). Top differentially expressed genes (DEG) were identified with a cubic spline regression model. Gene modules from normalized DEG expressions have been determined using a Dirichlet Process Gaussian Process (DPGP) mixture model. DEG list have been used for the detection of key TRs using REGulator-Gene Association Enrichment (REGGAE) and DbTFs among the TRs using TFcheckpoint databases. Thereafter, we used key TRs’ information in each of the gene modules to find top clusters by ranking with the number of key TRs inside them. We have labeled the top modules as “key modules”. We have also reconstructed GAN of DEGs with the help of a regularized dynamic partial correlation approach to find top genes in the GAN by ranking them according to their centrality measures in the network. We, then, predicted gene functions of the top genes using the GeneMANIA web server and analyzed obtained genes for their association with PDAC. We identified 8 TR’s and 10 genes in the reconstructed GAN to be directly associated with PDAC. We have detected 24 directly associated genes with PDAC in the top 4 key modules. Our analysis also reveals that only 24 out of 32 key gene modules contained all the key TRs. KEGG pathway, biological processes GO terms and DisGeNet analysis of top key modules were also determined.

Additionally, our analysis can be further extended by combining several biologically significant data sources for this disease by data integration techniques. Common symptoms of PDAC include weight loss, indigestion, abdominal and back pain. Thus, studying network pathways of the key gene modules may help unravel the serious complexities of this disease.

Moreover, the genes in the key modules may be further validated using *in vitro* experiments to reveal some important findings in pancreatic ductal adenocarcinoma and it’s pathogenesis. One may also verify the role of key regulators in the modules being potential biomarkers. Survival analysis using key transcriptional regulators and Protein-Protein interaction networks (PPIN) may enlighten us with greater insights about this disease.

## Acknowledgements (not compulsory)

Acknowledgements should be brief, and should not include thanks to anonymous referees and editors, or effusive comments. Grant or contribution numbers may be acknowledged.

## Author contributions statement

Must include all authors, identified by initials, for example: A.A. conceived the experiment(s), A.A. and B.A. conducted the experiment(s), C.A. and D.A. analysed the results. All authors reviewed the manuscript.

## Additional information

To include, in this order: **Accession codes** (where applicable); **Competing interests** (mandatory statement).

The corresponding author is responsible for submitting a [competing interests](https://www.nature.com/srep/policies/index.html#competing) statement on behalf of all authors of the paper.This statement must be included in the submitted article file.

