## Supplementary File 1 for "Discovering Key Transcriptomic Regulators in Pancreatic Ductal Adenocarcinoma using Dirichlet Process Gaussian Mixture Model"

Supplementary Figure 1: Reconstructed Gene Association Network using splineNetRecon with top 150 edges based on partial correlation score with Betweenness Centrality scores.

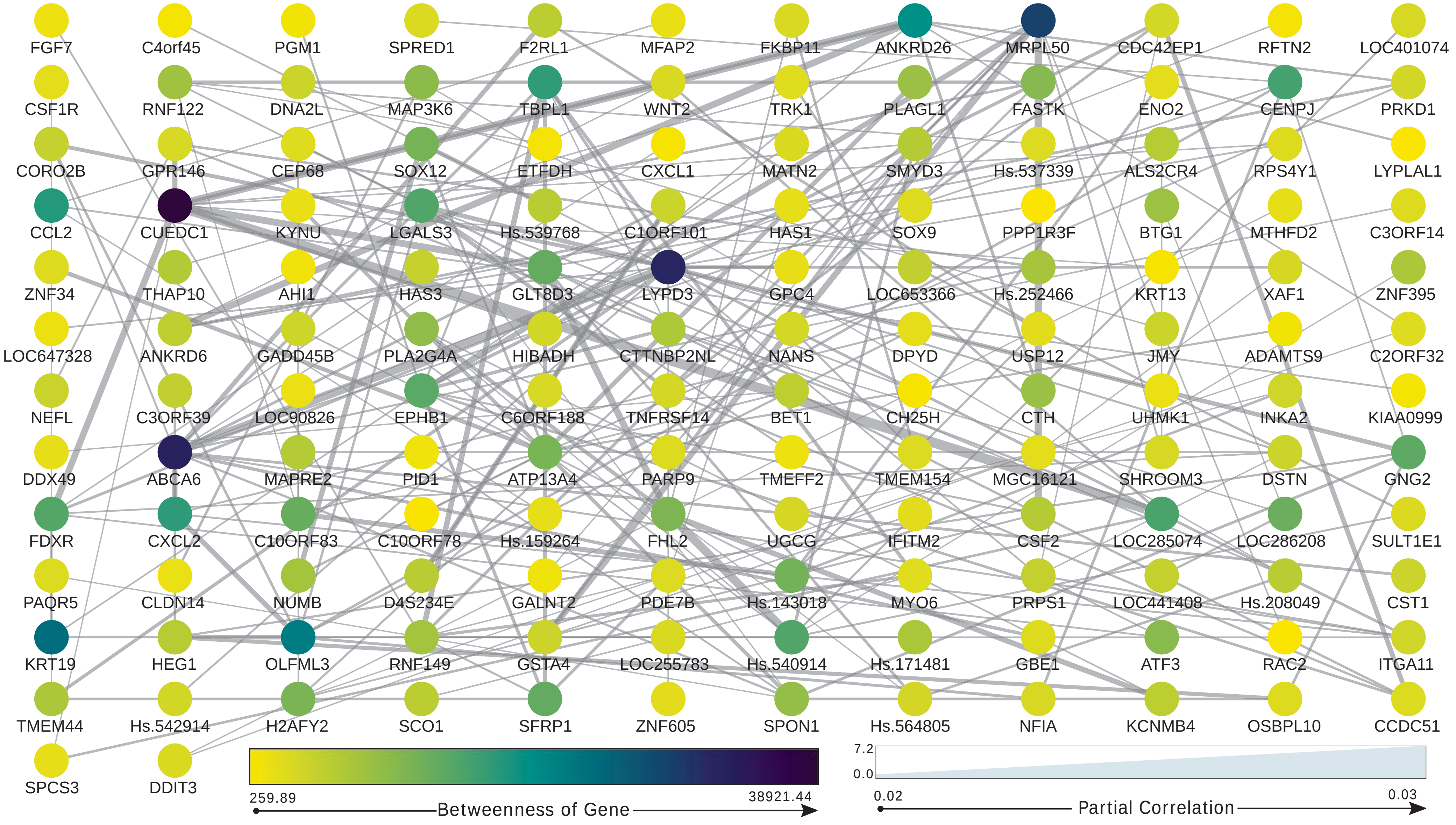

Supplementary Figure 2: Reconstructed Gene Association Network using splineNetRecon with top 150 edges based on partial correlation score with Closeness Centrality scores.

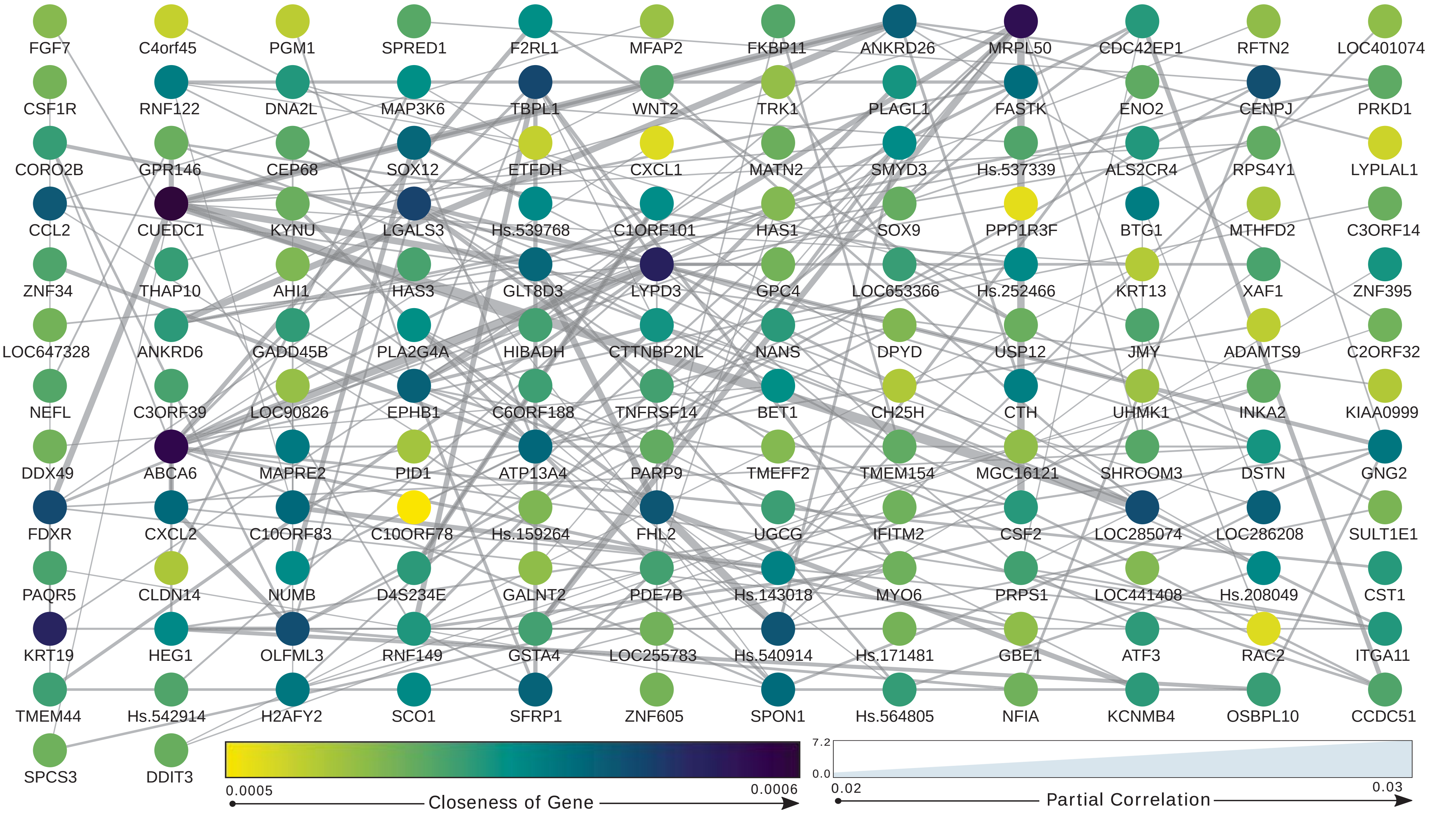

Supplementary Figure 3: A. KEGG Pathway of Key Module 2, B. Biological Process of Key Module 2, C. DisGeNet analysis of Key Module 2.

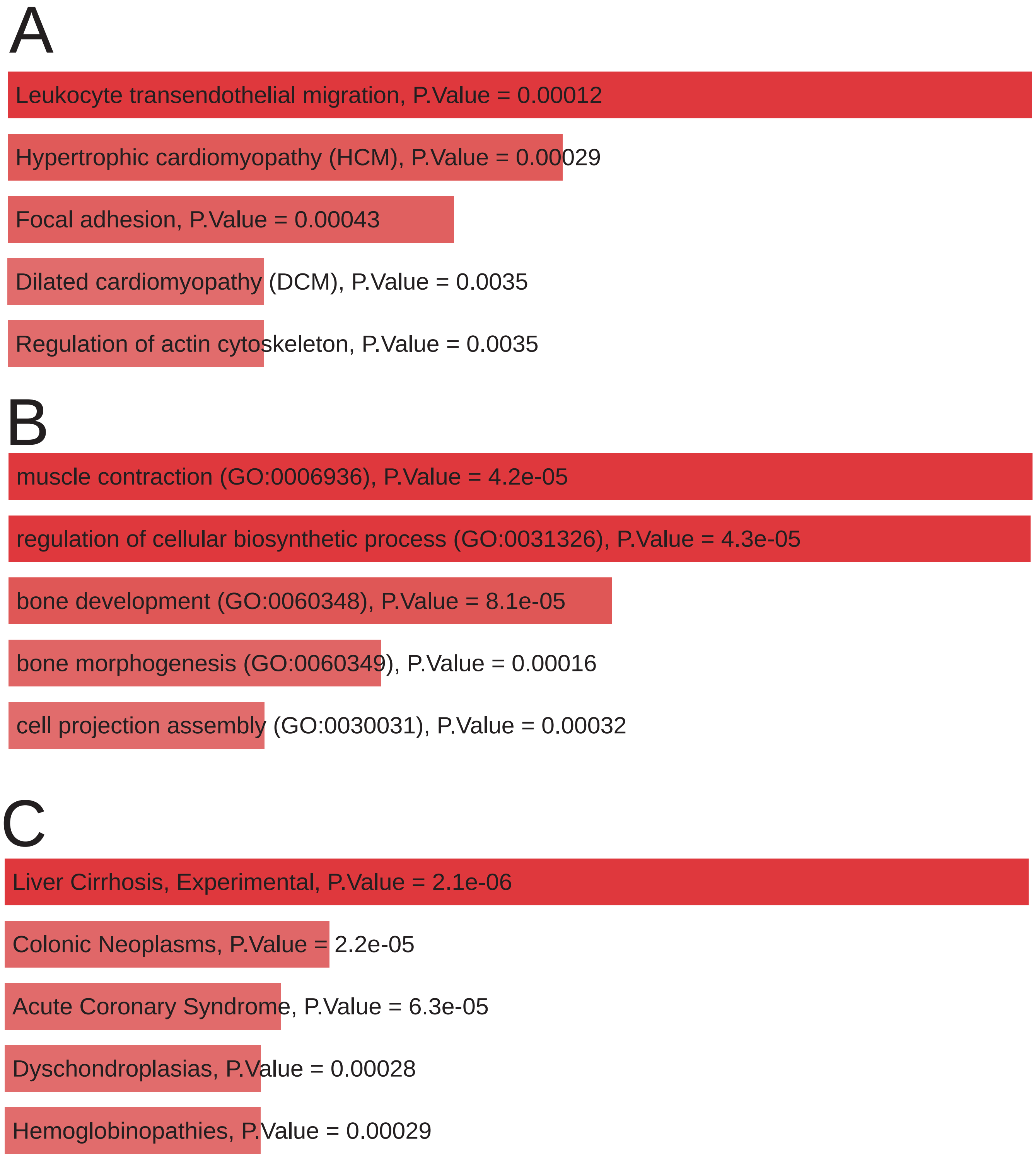

Supplementary Figure 4: A. KEGG Pathway of Key Module 3, B. Biological Process of Key Module 3, C. DisGeNet analysis of Key Module 3.

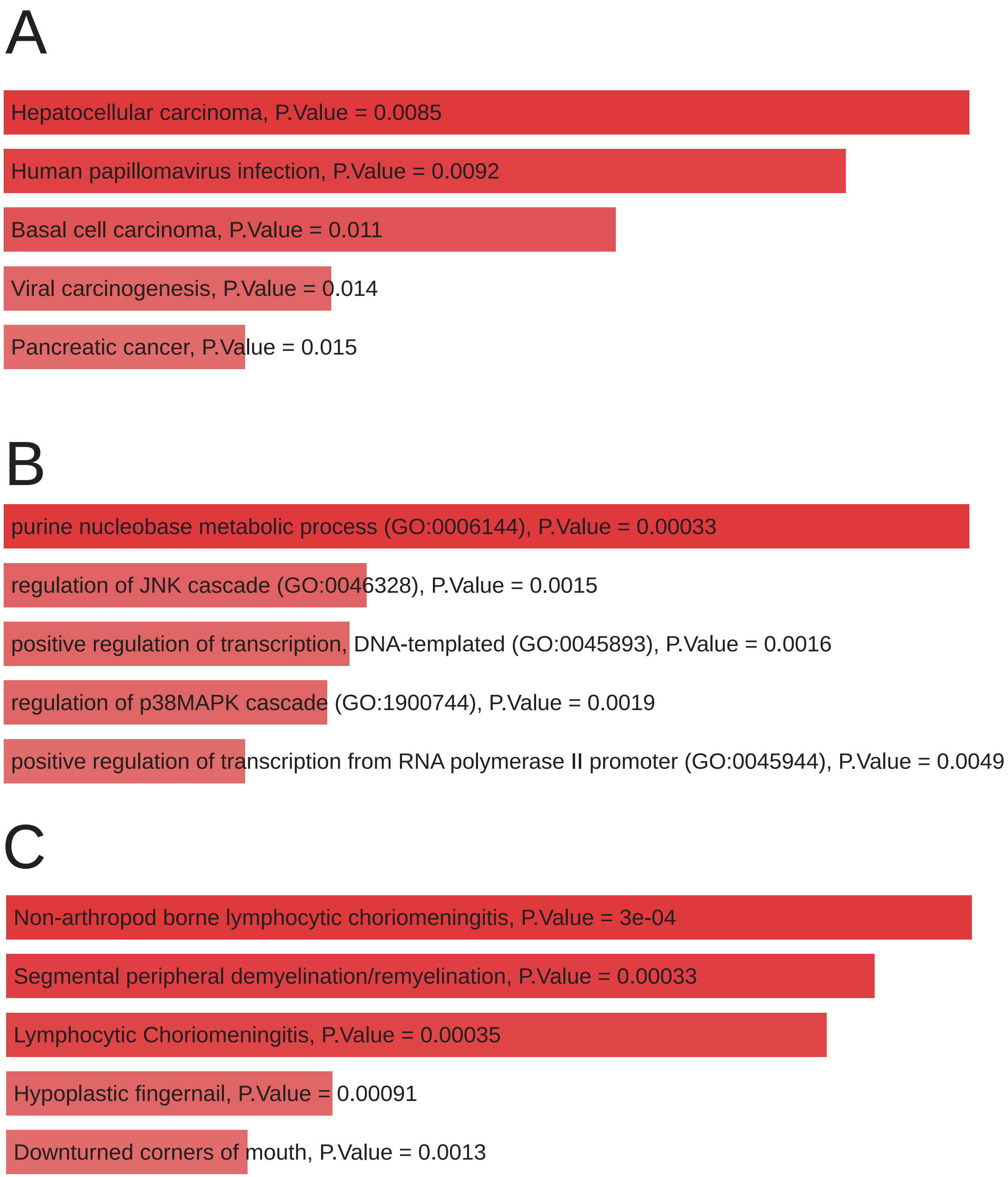

Supplementary Figure 5: A. KEGG Pathway of Key Module 4, B. Biological Process of Key Module 4, C. DisGeNet analysis of Key Module 4.

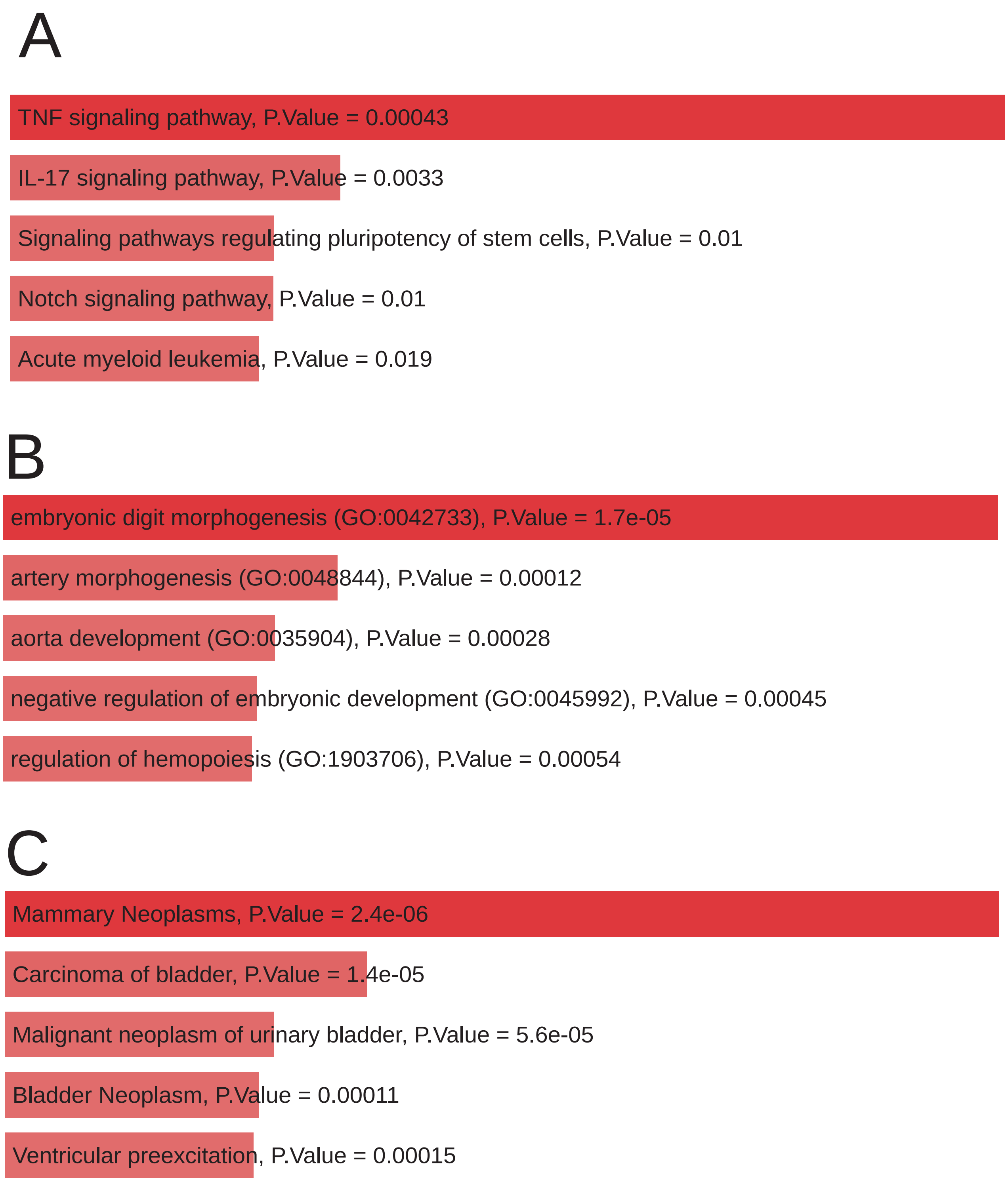

Supplementary Figure 6: Lollipop plot describing percentage of genes in key module 2 contributing to top 15 Gene Ontology terms.

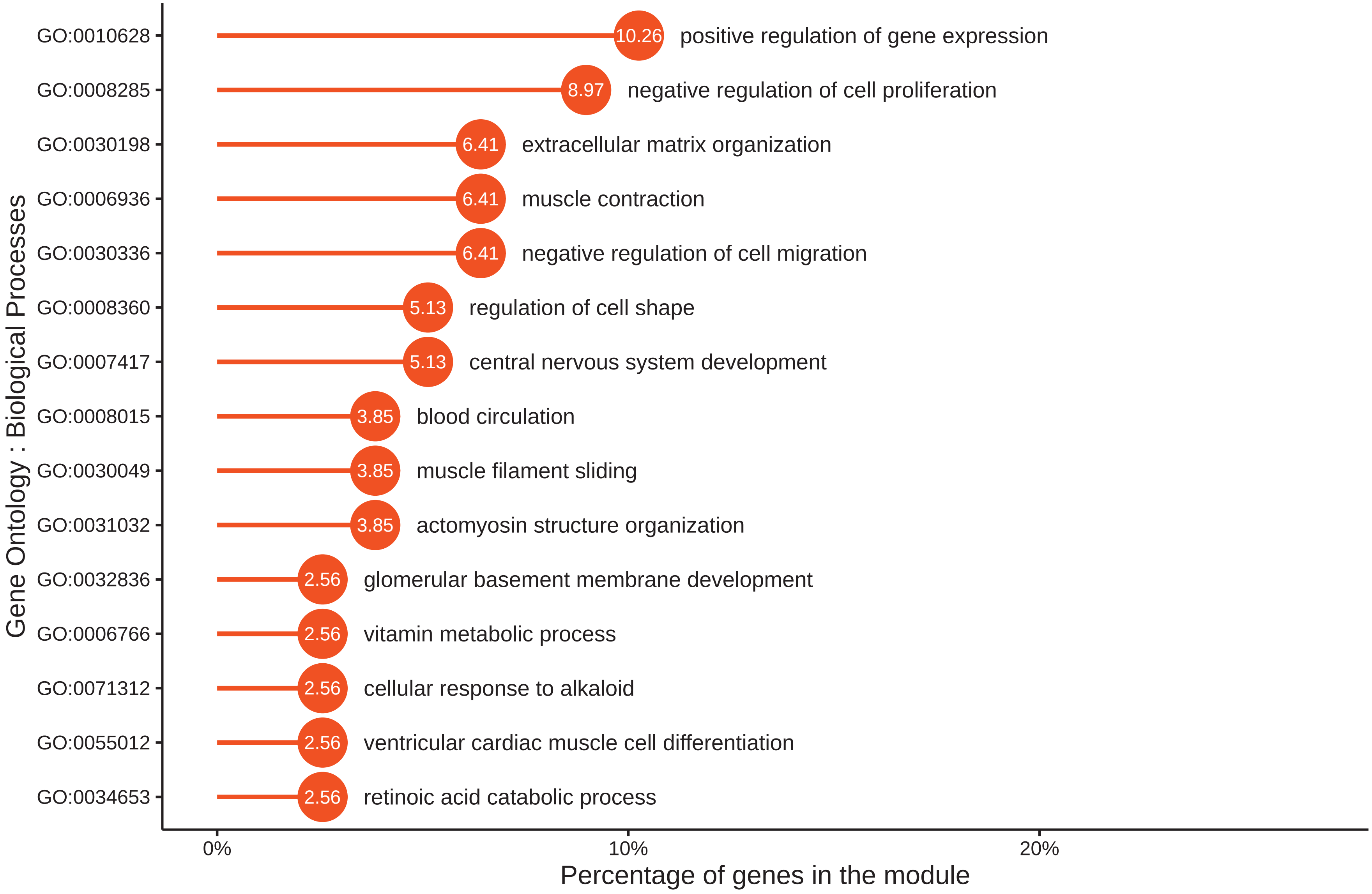

Supplementary Figure 7: Lollipop plot describing percentage of gene in key module 3 contributing to top 15 Gene Ontology terms.

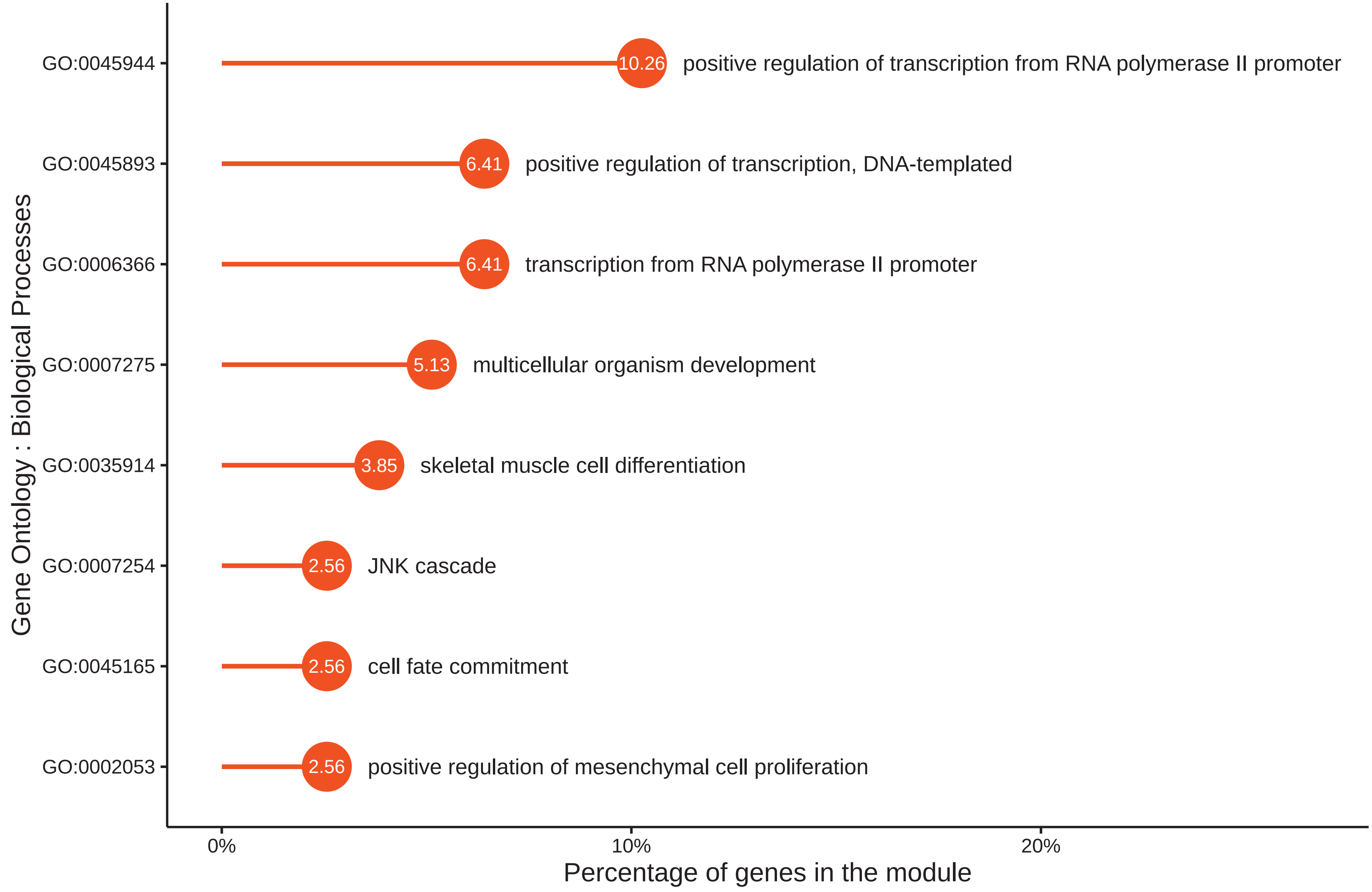

### Supplementary Figure 8: Lollipop plot describing percentage of gene in key module 4 contributing to top 15 Gene Ontology terms.

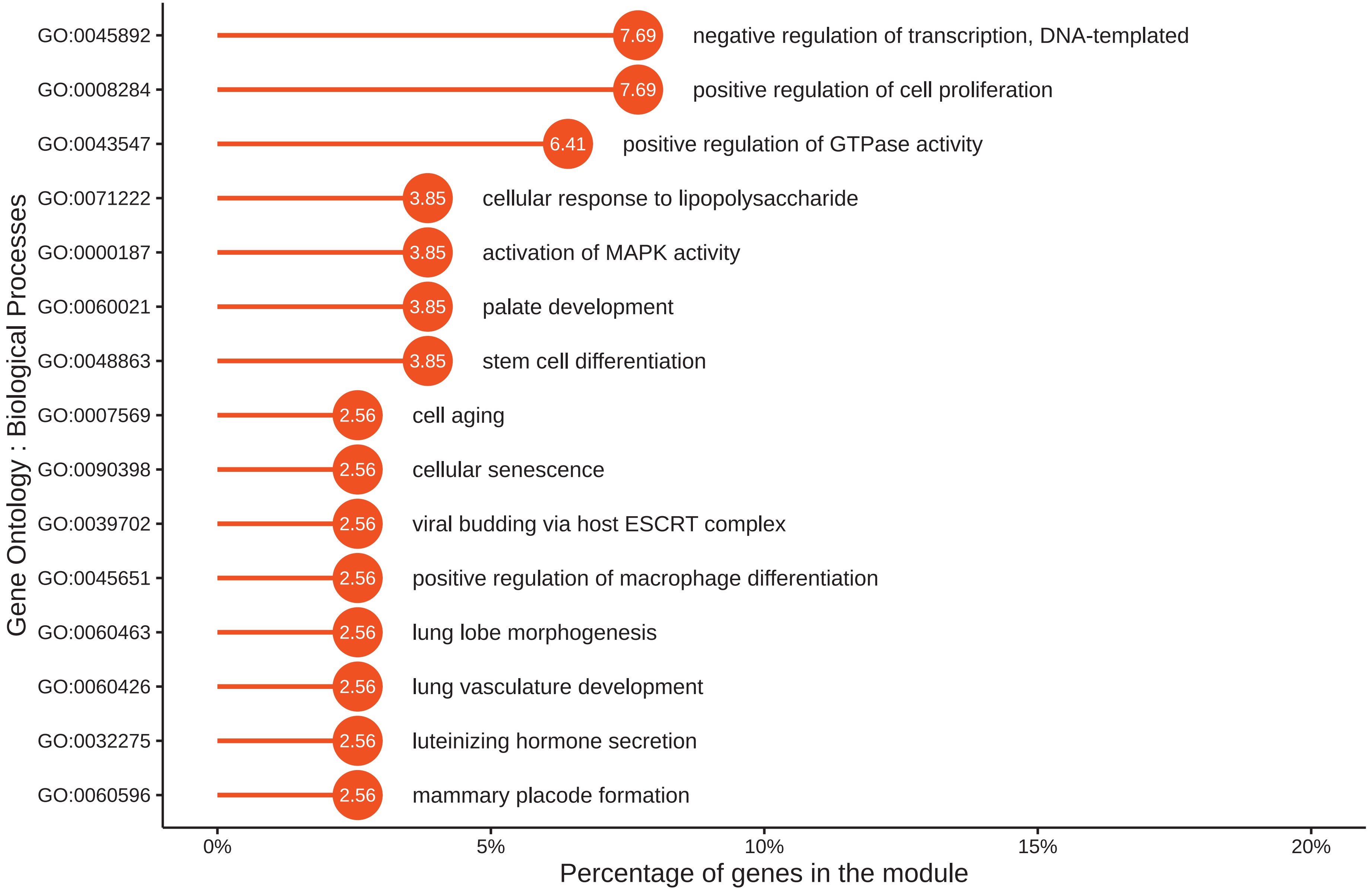
